# Contrasting taxonomic and functional diversity of tropical anuran communities across crude oil-contaminated and reference streams in a biodiversity hotspot

**DOI:** 10.64898/2026.11.25.769224

**Authors:** Mileidy Betancourth-Cundar, María Paula Toro-Gómez, Juan Camilo Ríos-Orjuela, Cristian Flórez Pai, María Victoria Posada-Vera, Lauren O’Connell

## Abstract

Conservation assessments of chronic anthropogenic disturbance increasingly ask whether changes in species composition translate into comparable changes in ecological function. This distinction is central to anticipating the long-term consequences of disturbance for ecosystem processes. Crude oil spills are a recurrent but understudied source of chronic disturbance in tropical freshwater ecosystems. We evaluated whether chronic oil contamination is associated with changes in taxonomic and functional diversity of anuran communities in six streams of the Nariño coastal piedmont (Colombian Chocó), within the area of influence of the Trans-Andean Pipeline. Amphibians are the most threatened vertebrate group worldwide and are particularly sensitive to both aquatic and terrestrial habitat degradation, making them sensitive indicators of chronic disturbance. We compared three oil-contaminated and three reference streams using standardized nocturnal surveys, Hill-number diversity, beta diversity, and multidimensional functional traits. We recorded 169 individuals from 24 species. Reference streams collectively contained more species and individuals than oil-contaminated streams, although coverage-standardized diversity varied substantially among individual streams and confidence intervals broadly overlapped. In contrast, functional diversity metrics, including functional richness, evenness, divergence, and trait composition, showed broad overlap between stream types. Overall, taxonomic patterns tended to favor reference streams, but these differences were not accompanied by a clear functional simplification, revealing a partial decoupling between taxonomic and functional responses consistent with high functional redundancy in one of the world’s most biodiverse and chronically oil-exposed tropical regions. This decoupling underscores the need to incorporate functional, not only taxonomic, metrics into biodiversity impact assessments of chronic contamination in biodiversity hotspots.

## Introduction

Crude oil spills are among the most severe anthropogenic disturbances to aquatic and riparian ecosystems, with effects ranging from acute mortality to persistent alterations in community structure, biogeochemical cycles, and ecosystem functioning (Lee, 2011; Chang et al., 2014; Perhar & Arhonditsis, 2014). Although crude oil spills cause immediate and widespread mortality riparian and aquatic-dependent biota at affected sites (Barron et al., 2020), their medium- and long-term ecological consequences remain poorly understood, particularly in species-rich tropical regions where environmental heterogeneity, species turnover, and high regional biodiversity may modulate ecological responses and recovery following disturbance (López-Delgado et al., 2020; Agra et al., 2021). Colombia is one of Latin America’s leading oil producers, and much of its oil infrastructure runs through ecologically sensitive regions (Dunning & Wirpsa, 2004; Guerrero-Useda, 2021). An emblematic case is the Trans-Andean Pipeline (hereafter “OTA”), which connects oil-producing fields in southern Colombia to the Pacific coast, spanning approximately 307 km. This pipeline crosses rainforests, agricultural areas, rivers, and the territories of Indigenous, peasant, and Afro-descendant communities whose livelihoods depend directly on the integrity of aquatic and terrestrial ecosystems. In this context, spills associated with the OTA represent a recurring but poorly understood threat to biodiversity and the socio-ecological security of local communities.

This situation is particularly significant in the Pacific piedmont of Nariño, a transitional zone between the Andes and the Pacific that forms part of the Tumbes–Chocó–Magdalena Biodiversity Hotspot, a globally recognized center of exceptional species richness, endemism, and environmental heterogeneity (Myers et al., 2000; Mittermeier et al., 2011; Pérez-Escobar et al., 2019). The region supports extraordinary biodiversity across multiple taxonomic groups, including plants, amphibians, reptiles, and birds (Castaño-Mora et al., 2004; Lynch & Suárez-Mayorga, 2004; Rangel, 2004; Flórez-Paí, 2016). Despite its ecological importance, it is increasingly threatened by multiple anthropogenic pressures, including deforestation, infrastructure expansion, extractive industries, and armed conflict (Hanson et al., 2009; González et al., 2018; Álvarez et al., 2021; Hanson, 2018). Crude oil contamination therefore represents an additional stressor that may interact with these ongoing disturbances, further compromising ecosystem integrity and the resilience of biological communities.

In the Pacific piedmont of Nariño, the OTA’s infrastructure has historically been the target of armed attacks, sabotage, and illegal oil extraction by non-state armed groups, leading to chronic and spatially heterogeneous contamination of water bodies and riparian forests (Miranda & Restrepo, 2005; Guerrero-Useda, 2021). One of the most serious incidents occurred in 2015, when a spill of nearly 14,000 barrels of crude oil contaminated the lower basins of the Mira and Rosario rivers, affected approximately 111 km of tropical rainforest riverbanks, and deprived more than 160,000 residents of Tumaco of access to drinking water (Garcés-Ordóñez & Espinosa, 2019; Ríos-Mármol et al., 2023). However, beyond these massive events, the installation of illegal valves, clandestine crude oil processing, localized leaks, and the dismantling of makeshift oil-processing sites by authorities collectively result in recurrent and multi-pathway contamination that has rarely been assessed from a community ecology perspective.

Amphibians are a particularly suitable group for assessing this type of disturbance. Their high skin permeability, dependence on both aquatic and terrestrial habitats throughout their life cycle, physiological sensitivity to microenvironmental changes, and central role in food webs make them excellent bioindicators of ecosystem health (Duellman & Trueb, 1986; Hopkins, 2007; Becker et al., 2023). Moreover, amphibians are currently the most threatened group of vertebrates on the planet: nearly 43% of species are listed as threatened under at least one threat category or have shown sustained population declines over the past four decades (Shaffer et al., 2015; Powers & Jetz, 2019; Luedtke et al., 2023). In this context, assessing amphibian responses to oil spills not only allows for a better understanding of the ecological impacts of this disturbance but also helps anticipate broader changes in ecosystem structure and functioning. This vulnerability is especially relevant in the Chocó Biogeographic Region, where amphibians exhibit exceptional levels of diversity and endemism, making them a critical indicator group for evaluating the ecological consequences of oil spills in one of the world’s most biodiverse regions.

Studies examining the effects of petroleum contamination on amphibians have focused on acute toxicity, larval mortality, physiological responses in laboratory settings, or individual-level effects (Mahaney, 1994; Zhang et al., 2022; Awkerman et al., 2024), while community-level and functional consequences remain largely unexplored, particularly in tropical systems. This functional perspective is particularly informative because it allows assessment not only of species loss, but also of whether ecological strategies represented within communities are altered and whether changes in functional redundancy may reduce ecosystem resilience to future disturbances (Mason et al., 2005; Villéger et al., 2008; Laliberté & Legendre, 2010; Mori et al., 2013; Ricotta et al., 2016).

In this study, we assess whether streams classified according to evidence of historical oil contamination differ from reference streams in the taxonomic richness and functional structure of amphibian communities in the Pacific piedmont of Nariño, within the area of influence of the Trans-Andean Pipeline. We evaluated whether contamination status was associated with differences at multiple levels of community organization: (i) differences in taxonomic richness and relative abundance; (ii) changes in the extent of functional space; (iii) functional homogenization of morphological traits; and (iv) alterations in the representation of ecological strategies.

Under the hypothesis that environmental conditions associated with oil contamination can act as a filter on species sensitive to riparian microhabitats, we predicted that streams classified as oil-contaminated would exhibit lower taxonomic richness, a contracted functional space, and shifts in the distribution of traits associated with locomotion, microhabitat use, and behavior. To evaluate these predictions, we integrated alpha and beta diversity metrics, classic indices of functional diversity, moments of trait distributions (such as mean, variance, skewness, and kurtosis), and multivariate functional space analyses in a tropical system subjected to chronic contamination within one of the planet’s most important biodiversity hotspots.

## Materials and Methods

### Study site selection

To select study sites, we reviewed local environmental records of oil spills associated with the OTA in the Altaquer–El Diviso corridor (Barbacoas, Nariño, Colombia) (Fig. 1). With the assistance of local guides, we surveyed streams within three nature reserves: the Río Ñambí Nature Reserve, the Bangsias Bird Lodge Nature Reserve, and the Inkal Awá La Nutria Piman Nature Reserve (Fig. 1A). For each stream, we recorded geographic coordinates and, integrating visual inspection with traditional ecological knowledge, assessed whether streams were contaminated by petroleum. Streams were classified as uncontaminated (reference) (Fig. 1B) and contaminated based on visible signs of crude oil contamination residues (e.g., sheen presence, odor, and stained sediments) on the water surface, substrate rocks, riparian soils, and vegetation along the riverbanks (Fig. 1C).

**Fig. 1.**
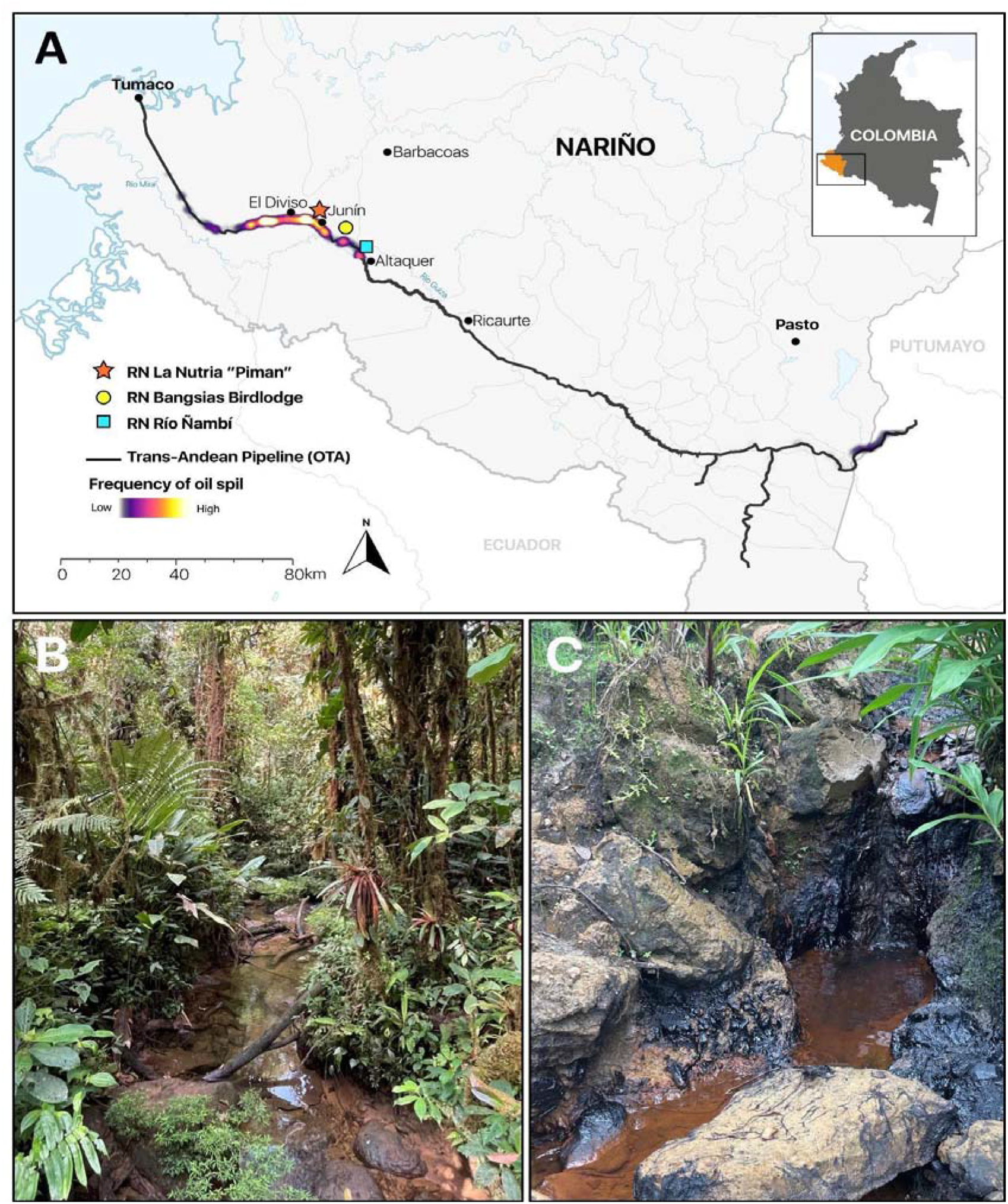
Oil contamination in the study region is concentrated along the trans-Andean pipeline corridor and overlaps part of the sampled frog habitats in Barbacoas, Nariño. (a) Location of the six sampled streams in the department of Nariño, Colombia, relative to the Trans-Andean Oil Pipeline (Oleoducto Transandino, OTA). The color gradient along the pipeline represents the historical frequency of oil spill events, from low to high incidence. The map also shows the three protected areas included in the study (Inkal Awá La Nutria “Piman” Natural Reserve, Bangsias Birdlodge Reserve, and Río Ñambí Natural Reserve), the pipeline route, major rivers, and populated areas. The inset indicates the location of the study region within Colombia. (b) Representative reference stream located within well-preserved forest. (c) Representative oil-contaminated stream showing petroleum residues on the streambed. Map lines delineate the study area and do not necessarily depict accepted national, provincial, or other boundaries

Because historical oil spills are spatially restricted, contaminated streams could not be randomly assigned. Therefore, for each oil-contaminated stream we selected a reference stream with similar environmental characteristics (e.g. elevation, stream size), while lacking visible evidence of oil contamination (Table S1). This matching approach was intended to minimize environmental differences unrelated to contamination. Based on these criteria, we selected six streams: three uncontaminated (reference) and three oil-contaminated. The reference streams were Gualgaro II (Río Ñambí NR), El Acueducto (Bangsias Bird Lodge NR), and La Nutria (Inkal Awá La Nutria Piman NR). The streams classified as oil-contaminated were El Lindero (Bangsias Bird Lodge NR), El Tejón, and Río Caraño (Inkal Awá La Nutria Piman NR). We conducted three field expeditions of ten days each between March and September 2024, spanning the dry and wet seasons. Each stream was surveyed at least four times, with 3–7 observers participating in each survey. Across the six streams, total sampling effort was 242.53 person-hours. Nocturnal visual encounter surveys were conducted between 17:00 and 21:00 h along 100-m transects covering the stream edge and a parallel transect within the riparian forest, 3–5 m from the stream channel (Jaeger, 1994; Rueda et al., 2006). Because sampling completeness varied among streams, taxonomic diversity comparisons were subsequently standardized by sample coverage rather than assuming equivalent sampling completeness across sites.

### Ecological and morphological data collection

We identified all detected individuals to the finest possible taxonomic level, recording activity, observation substrate, and perch height for each individual (Heyer et al., 1994). We photographed each individual from dorsal and ventral views with a metric reference scale. We released all individuals at the capture site and handled them with sterile gloves throughout. Species were identified in the field using photographic records and biological archives maintained by the local reserves, as no voucher specimens were collected. Individuals that could not be identified to species level were treated as distinct morphospecies (*Pristimantis* sp. 1–7) and included in all taxonomic and functional analyses. Conservation status was assigned following the IUCN Red List of Threatened Species (IUCN, 2024).

To characterize the functional dimensions of the communities, we measured six continuous morphological traits directly in the field: body size measured as snout-vent length (SVL), mouth width (MW), forearm length (FA), femur length (FL), and tibiofibula length (TF). We measured body mass using a precision digital scale (0.01 g), controlling for water retention before each measurement (Meuche et al., 2012). We also recorded categorical ecological attributes, including microhabitat use (arboreal or terrestrial), observation substrate, activity period (diurnal or nocturnal), behavioral state at observation (perched or resting), and degree of interdigital foot webbing (absent, basal, moderate, extensive, or fully webbed). Because surveys were conducted exclusively between 17:00 and 21:00 h, activity period and behavioral state were strongly conditioned by the nocturnal sampling protocol and were therefore retained only as descriptive variables. Comparisons of categorical functional composition focused on microhabitat use, substrate type, and foot webbing, which were less directly constrained by survey timing. These traits capture complementary dimensions of locomotion and microhabitat use (Salgado Negret et al., 2016; Blanco-Torres et al., 2019).

All data were collected in accordance with applicable ethical standards and animal-use regulations, with the prior authorization and accompaniment of the Indigenous communities involved throughout the research process. Biological sampling was conducted under the Framework Collection Permit granted to Universidad de los Andes by Resolution No. 1177 of 9 October 2014 (IDB 0359).

### Taxonomic alpha and beta diversity

To assess differences in amphibian anuran diversity between oil-contaminated and reference streams, we estimated taxonomic alpha and beta diversity. We quantified alpha diversity using a coverage-based rarefaction and extrapolation framework, which standardize comparisons among sites with unequal sampling completeness effort (Chao & Jost, 2012; Chao et al., 2014). We estimated diversity, as Hill numbers for orders q = 0, 1, and 2, corresponding to species richness, the exponential of Shannon entropy, and the inverse Simpson index, respectively (Jost, 2007), using the iNEXT package v. 3.0.2 (Hsieh et al., 2025) in R v.4.5.1 (R Core Team, 2025). We analyzed abundance data and estimated 95% confidence intervals from 200 bootstrap replicates.

Because sample coverage differed among streams, we standardized diversity comparisons to a common sample coverage of 0.70, selected conservatively below the lowest observed coverage (0.735) so that all assemblages were compared by rarefaction at an equivalent level of sampling completeness. To assess taxonomic turnover between sites, we calculated pairwise compositional similarity using the abundance-based Chao-Jaccard index and estimated beta diversity corrected for detectability and abundance of rare species following Jost (2007). At the regional scale, we partitioned diversity into alpha, gamma, and multiplicative beta diversity using coverage-standardized Hill numbers following Jost (2007), allowing us to estimate the effective number of compositionally distinct communities across the study landscape.

### Functional diversity

To assess whether oil contamination is associated with changes in the functional structure of amphibian communities, we analyzed functional diversity across six streams (three contaminated and three reference) by integrating metrics of functional range, trait distribution, and ecological composition. This integrative approach enabled detection of shifts in functional space occupation and potential signals of environmental filtering on the ecological strategies represented within communities (Villéger et al., 2008; Laliberté & Legendre, 2010; Mouillot et al., 2013).

We calculated classic functional diversity indices (FRic, FEve, FDiv, and FDis) based on a species × site abundance matrix and a matrix of continuous traits including body mass, snout-vent length (SVL), mouth width (MW), forearm length (FA), femur length, tibiofibula length, and perch height. This trait was quantified as the vertical distance from the ground to the observation substrate, with species-level means used in all functional analyses. This variable captures vertical microhabitat partitioning, a functionally relevant axis associated with thermoregulation and desiccation avoidance in tropical anurans (Scheffers et al., 2014; Nowakowski et al., 2018). Although previous frameworks have treated vertical habitat use categorically in Colombian amphibians (Blanco-Torres et al., 2019), continuous quantification improves functional resolution by reducing the information loss inherent to discrete categorization (Violle et al., 2007). Together, these traits capture the primary axes of functional variation shaping amphibian communities (Salgado Negret et al., 2016; Blanco-Torres et al., 2019). Continuous traits were log(x + 1)-transformed and standardized to zero mean and unit variance prior to analysis. Functional diversity indices were calculated using dbFD in the FD package (Laliberté et al., 2014), with stand.x = FALSE because traits had been standardized beforehand, ord = "metric", corr = "cailliez" to correct for negative eigenvalues, and m = "max" (calc.FRic = TRUE, calc.FDiv = TRUE; Legendre & Legendre, 2012). Additionally, we regressed FRic against species richness to obtain residual functional richness (FRic_resid), isolating functional changes independent of differences in taxonomic richness.

To assess shifts in the shape of trait distributions within communities, we calculated the community-weighted trait means (CWM), weighted standard deviation (CWsd), skewness, and kurtosis for each site and trait. This approach enabled detection of potential signals of environmental filtering and functional homogenization that may not be evident from global functional diversity indices alone (Spasojevic & Suding, 2012; Mason et al., 2013). CWM and CWsd were calculated using the Hmisc package, and skewness and kurtosis using the moments package in R (Harrel & Dupont, 2024; Komsta & Novomestky, 2022).

To assess the multivariate configuration of functional space, we used the log (x + 1)-transformed and standardized continuous trait matrix to calculate community-level functional centroids, global functional dispersion, and centroid displacement. For each community, the functional centroid was calculated as the abundance-weighted mean position of species in the full standardized trait space. Global functional dispersion was calculated as the abundance-weighted mean Euclidean distance of species to their community centroid, and centroid displacement was calculated as the Euclidean distance between each community centroid and the overall regional centroid. Principal component analysis (PCA) was used only to visualize the multivariate functional space by projecting community centroids onto the PC1–PC2 and PC1–PC3 planes; PC1 and PC2 jointly explained 90.1% of total trait variation, and the variance explained by each individual axis is reported in the results section. These metrics allowed us to evaluate shifts in the position and structure of functional space between oil-contaminated and reference streams (Mouillot et al., 2013). Finally, to assess shifts in categorical ecological composition, we calculated abundance-weighted proportions of microhabitat use, substrate type, and interdigital foot webbing for each stream. Activity period and behavioral state were summarized descriptively but excluded from between-condition inference because their observed distributions were strongly constrained by the nocturnal sampling protocol. For the focal categorical traits, we calculated the proportional representation of each category at each site and compared these proportions between contamination conditions using the exact permutation framework described below.

Given the small number of sites, we compared functional metrics between oil-contaminated and reference streams using custom exact permutation tests implemented in R. The test statistics were the absolute difference in group means, and all 20 possible group allocations preserving the three-site sample size per condition were enumerated. P-values were calculated as the proportion of permuted differences equal to or greater than the observed difference. Because the statistics were based on the absolute difference between group means, complementary allocations yield identical test statistics, resulting in a minimum attainable two-sided p-value of 0.10. Accordingly, permutation p-values were interpreted descriptively rather than as definitive evidence for the presence or absence of differences between stream categories, and inference emphasized the direction, magnitude, overlap, and consistency of patterns among individual sites. We additionally performed leave-one-out sensitivity analyses, sequentially excluding sites from each condition, to assess whether the qualitative patterns depended strongly on particular streams; these analyses were interpreted as sensitivity checks rather than independent significance tests.

## Results

### Taxonomic alpha and beta diversity

Across the six stream sites, we recorded 169 anuran individuals belonging to 24 species (Table 1; Fig. 2). Strabomantidae was the richest family, accounting for 14 of the 24 species, including seven unidentified *Pristimantis* morphospecies (*Pristimantis* spp. 1–7). Centrolenidae comprises four species, Hylidae three, Bufonidae two, Dendrobatidae and Ranidae one species. Two species were classified as Vulnerable (VU) (*Rhaebo andinophrynoides* and *Pristimantis laticlavius*), whereas *Oophaga sylvatica*, *Cochranella mache*, and *Pristimantis verecundus* were classified as Near Threatened (NT). *Pristimantis laticlavius* occurred in all six streams, whereas *Rhaebo andinophrynoides* was restricted to La Nutria. Reference streams contained 19 species and 95 individuals, whereas oil-contaminated streams contained 16 species and 74 individuals. We recorded 13 species in more than one stream and 11 species shared between reference and oil-contaminated streams. Eight species were exclusive to reference streams, whereas five occurred only in oil-contaminated streams. Species richness ranged from 6 to 12 species per stream. La Nutria had the highest richness, whereas El Acueducto and El Lindero had the lowest. Abundance ranged from 18 to 43 individuals, with the highest abundance in Gualgaro II and the lowest in El Acueducto and El Tejón.

**Fig. 2.**
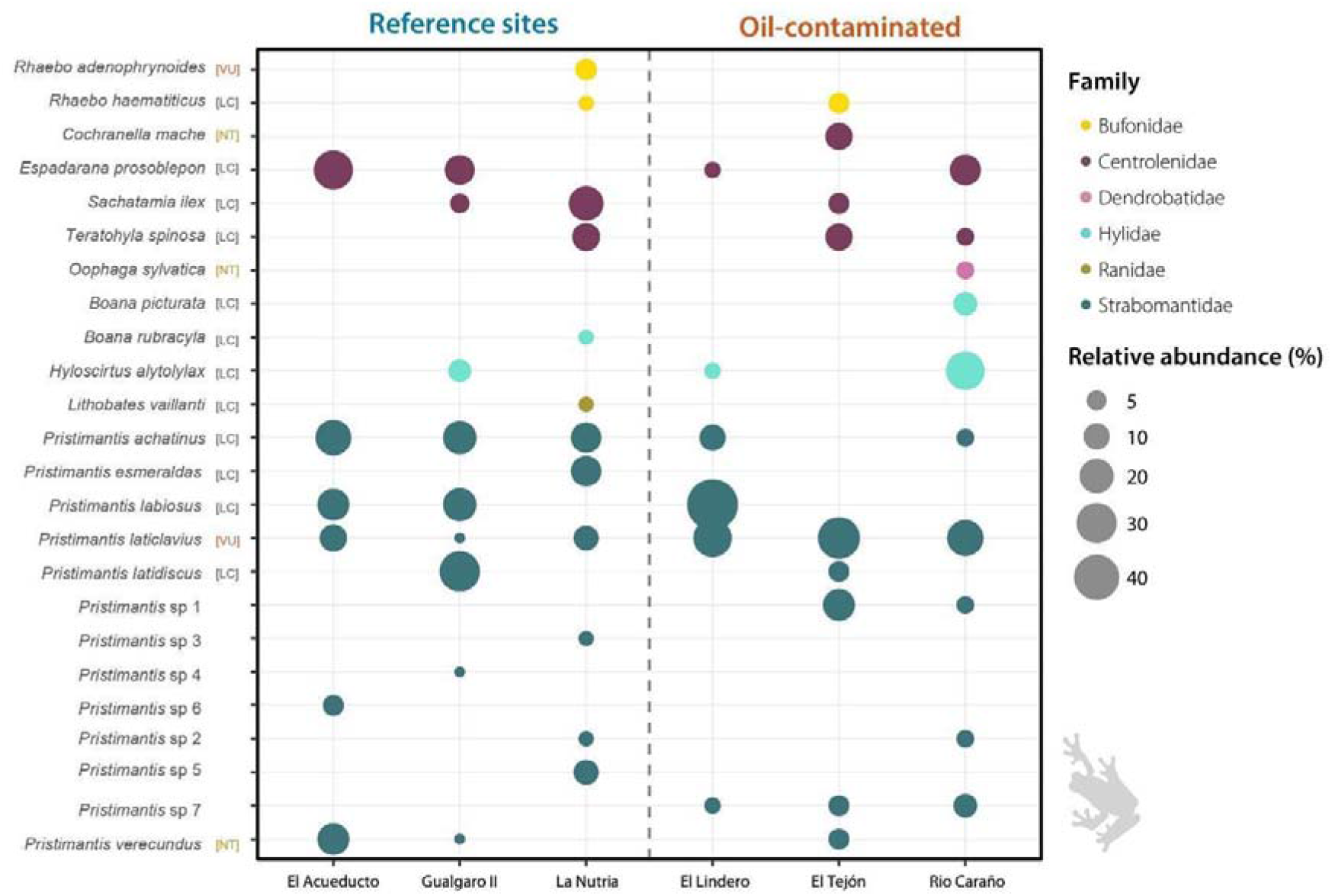
Relative abundance and taxonomic composition of anuran assemblages in reference and oil-contaminated streams. Bubble plot showing the relative abundance (%) of each anuran species recorded in the six sampled streams. Bubble size is proportional to the relative abundance of each species within each stream, and colors indicate taxonomic family. Reference streams (El Acueducto, Gualgaro II, and La Nutria) are shown on the left, whereas oil-contaminated streams (El Lindero, El Tejón, and Río Caraño) are shown on the right. Conservation status according to the IUCN Red List is indicated next to each species name (VU = Vulnerable, NT = Near Threatened, LC = Least Concern)

**Table 1.** Frog assemblages showed strong site-level turnover, with many species restricted to single streams and only a few widespread taxa shared across both reference and oil-contaminated sites. Presence–absence matrix of anuran species recorded during standardized nocturnal surveys in six streams of the Barbacoas coastal piedmont, Nariño, Colombia. Rows list species grouped by family, and columns indicate sampling sites: three reference streams (La Nutria, Gualgaro II, El Acueducto) and three oil-contaminated streams (El Lindero, El Tejón, Río Caraño). This table summarizes the taxonomic composition underlying all diversity and functional analyses. Species such as *Pristimantis laticlavius* and *Pristimantis achatinus* were recorded across multiple streams, whereas many others were site-restricted, reflecting substantial local heterogeneity in assemblage composition. Because species turnover was moderate across all sites, this table should be interpreted primarily as evidence of spatially structured community variation rather than as a simple separation between contamination categories

|  |  |  | **Reference sites** |  |  | **Oil-contaminated sites** |  |  |
|---|---|---|---|---|---|---|---|---|
| **Family** | **Species** | **IUCN Cat.** | **La Nutria** | **Gualgaro II** | **El Acueducto** | **El Lindero** | **El Tejón** | **Río Caraño** |
| Bufonidae | *Rhaebo andinophrynoides* | VU | x |  |  |  |  |  |
|  | *Rhaebo haematiticus* | LC | x |  |  |  | x |  |
| Dendrobatidae | *Oophaga sylvatica* | NT |  |  |  |  |  | x |
| Centrolenidae | *Sachatamia ilex* | LC | x | x |  |  | x |  |
|  | *Teratohyla spinosa* | LC | x |  |  |  | x | x |
|  | *Cochranella mache* | NT |  |  |  |  | x |  |
|  | *Espadarana prosoblepon* | LC |  | x | x | x |  | x |
| Hylidae | *Boana picturata* | LC |  |  |  |  |  | x |
|  | *Boana rubracyla* | LC | x |  |  |  |  |  |
|  | *Hyloscirtus alytolylax* | LC |  | x |  | x |  | x |
| Ranidae | *Lithobates vaillanti* | LC | x |  |  |  |  |  |
| Strabomantidae | *Pristimantis achatinus* | LC | x | x | x | x |  | x |
|  | *Pristimantis esmeraldas* | LC | x |  |  |  |  |  |
|  | *Pristimantis labiosus* | LC |  | x | x | x |  |  |
|  | *Pristimantis laticlavius* | VU | x | x | x | x | x | x |
|  | *Pristimantis latidiscus* | LC |  | x |  |  | x |  |
|  | *Pristimantis verecundus* | NT |  | x | x |  | x |  |
|  | *Pristimantis* sp 1 |  |  |  |  |  | x |  |
|  | *Pristimantis* sp 2 |  |  |  |  |  |  | x |
|  | *Pristimantis* sp 3 |  | x |  |  |  |  |  |
|  | *Pristimantis* sp 4 |  |  | x |  |  |  |  |
|  | *Pristimantis* sp 5 |  | x |  |  |  |  |  |
|  | *Pristimantis* sp 6 |  |  |  | x |  |  |  |
|  | *Pristimantis* sp 7 |  |  |  |  | x | x | x |

Sampling coverage ranged from 0.735 to 0.950 (Fig. 3), indicating moderate to high inventory completeness (Fig. 3A). Coverage was highest in El Acueducto (0.950) and lowest in El Tejón (0.735). We standardized diversity estimates to a common sample coverage of 0.70 using the coverage-based rarefaction and extrapolation framework of Chao and Jost.

**Fig. 3.**
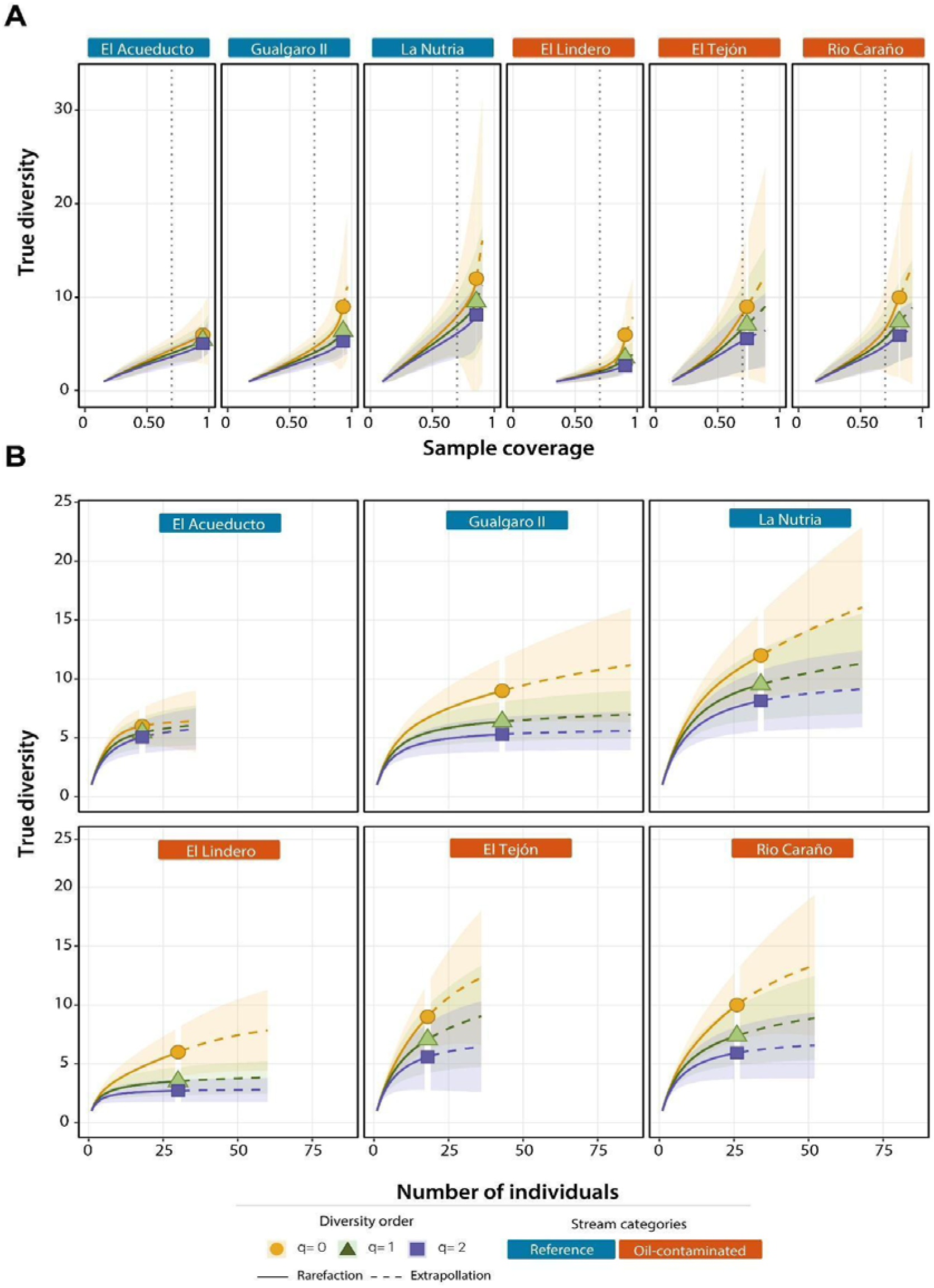
Oil-contaminated streams tended to support lower frog diversity than reference streams, especially for species richness and comparable sampling completeness. Hill-number rarefaction and extrapolation curves for frog assemblages in six streams in Barbacoas, Nariño. (a) Panels show taxonomic diversity estimated as species richness (q = 0), Shannon diversity (q = 1), and Simpson diversity (q = 2). In the upper panels, diversity is plotted as a function of sample coverage to standardize comparisons by inventory completeness (0.70); (b) in the lower panels, diversity is plotted as a function of the number of individuals sampled. Solid lines indicate interpolation (rarefaction), dashed lines indicate extrapolation, shaded bands represent 95% confidence intervals, and symbols indicate observed diversity values. Panel labels identify each stream, with white labels indicating reference streams and black labels indicating crude-oil polluted streams. These curves show that differences between stream types were strongest for richness (q = 0), whereas patterns for Shannon and Simpson diversity were more moderate

Coverage-standardized Hill numbers varied among streams (Fig. 3B; Table S2). La Nutria consistently showed the highest values of species richness (q = 0), Shannon diversity (q = 1), and Simpson diversity (q = 2), whereas El Lindero showed the lowest values. Reference streams generally exhibited higher diversity than oil-contaminated streams, although Río Caraño showed diversity values comparable to those of some reference streams. Confidence intervals overlapped broadly among streams, particularly for extrapolated estimates.

The abundance-based Chao–Jaccard index showed marked variation in species composition among streams, with pairwise similarity ranging from 0.16 to 0.74. Similarity was highest between El Acueducto and El Lindero (0.74) and lowest between El Tejón and Gualgaro II (0.16). Intermediate values were observed for Río Caraño-El Tejón (0.60), Gualgaro II-Río Caraño (0.59), Gualgaro II-El Acueducto (0.54), La Nutria-Río Caraño (0.44), and the remaining stream pairs (0.17-0.37).

Coverage-standardized Hill-number partitioning indicated moderate taxonomic differentiation among streams. Mean alpha diversity ranged from 4.24 effective species (q = 2) to 5.71 effective species (q = 0), whereas gamma diversity ranged from 7.11 to 9.86 effective species. Multiplicative beta diversity varied little among diversity orders, ranging from 1.68 to 1.73 effective communities (Table S2), indicating that the regional assemblage was equivalent to approximately 1.7 compositionally distinct effective communities.

### Functional diversity

Classic functional indices varied among sites, with FRic values close to zero in several streams and a maximum at the reference stream La Nutria (FRic = 0.281), which also exhibited the highest functional dispersion (FDis = 2.28; Table 2). Exact permutation p-values were 0.9 for FRic, 0.2 for FEve, 0.8 for FDiv, and 0.3 for FDis.

**Table 2.**
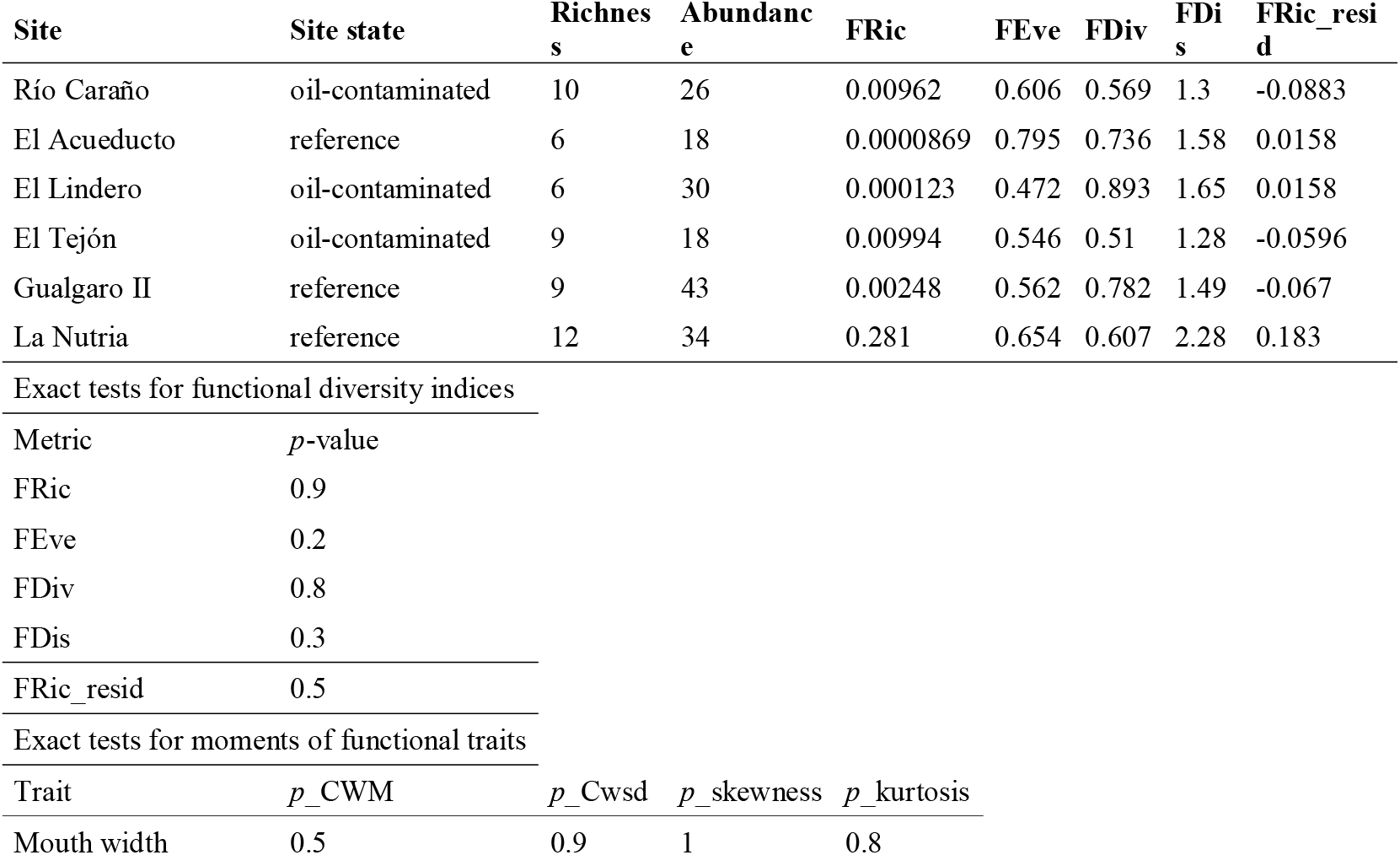
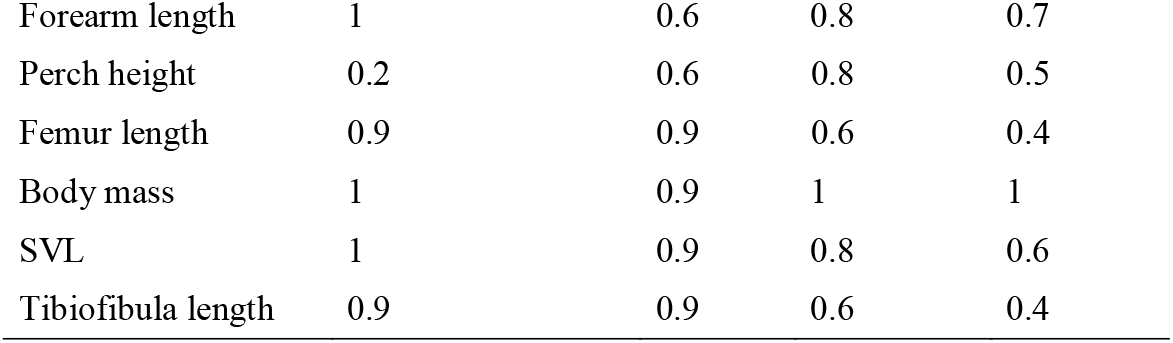
Functional diversity varied among streams, but no functional metric showed a consistent difference between oil-contaminated and reference sites. Stream-level taxonomic descriptors (species richness and abundance) and functional diversity indices for anuran assemblages sampled in six streams in Barbacoas, Nariño, Colombia. Functional metrics include functional richness (FRic), functional evenness (FEve), functional divergence (FDiv), functional dispersion (FDis), and richness-corrected residual functional richness (FRic_resid), which accounts for differences in richness among streams. Exact permutation p-values comparing stream categories (reference vs. oil-contaminated) are reported for each global functional diversity metric (FRic, FEve, FDiv, FDis). Because only three streams were available per condition, these p-values have limited inferential resolution and are presented together with the observed degree of overlap and among-stream variation. The table also reports exact test p-values for distributional moments of continuous traits, including community-weighted means (CWM), weighted standard deviations (CWsd), skewness, and kurtosis. Trait distributions also largely overlapped between stream types, indicating no clear evidence of directional trait filtering or functional homogenization associated with chronic oil contamination

| **Site** | **Site state** | **Richness** | **Abundance** | **FRic** | **FEve** | **FDiv** | **FDis** | **FRic_resid** |
|---|---|---|---|---|---|---|---|---|
| Río Caraño | oil-contaminated | 10 | 26 | 0.00962 | 0.606 | 0.569 | 1.3 | -0.0883 |
| El Acueducto | reference | 6 | 18 | 0.0000869 | 0.795 | 0.736 | 1.58 | 0.0158 |
| El Lindero | oil-contaminated | 6 | 30 | 0.000123 | 0.472 | 0.893 | 1.65 | 0.0158 |
| El Tejón | oil-contaminated | 9 | 18 | 0.00994 | 0.546 | 0.51 | 1.28 | -0.0596 |
| Gualgaro II | reference | 9 | 43 | 0.00248 | 0.562 | 0.782 | 1.49 | -0.067 |
| La Nutria | reference | 12 | 34 | 0.281 | 0.654 | 0.607 | 2.28 | 0.183 |
| Exact tests for functional diversity indices |  |  |  |  |  |  |  |  |
| Metric | *p*-value |  |  |  |  |  |  |  |
| FRic | 0.9 |  |  |  |  |  |  |  |
| FEve | 0.2 |  |  |  |  |  |  |  |
| FDiv | 0.8 |  |  |  |  |  |  |  |
| FDis | 0.3 |  |  |  |  |  |  |  |
| FRic_resid | 0.5 |  |  |  |  |  |  |  |
| Exact tests for moments of functional traits |  |  |  |  |  |  |  |  |
| Trait | *p*_CWM | *p*_Cwsd | *p*_skewness | *p*_kurtosis |  |  |  |  |
| Mouth width | 0.5 | 0.9 | 1 | 0.8 |  |  |  |  |
| Forearm length | 1 | 0.6 | 0.8 | 0.7 |  |  |  |  |
| Perch height | 0.2 | 0.6 | 0.8 | 0.5 |  |  |  |  |
| Femur length | 0.9 | 0.9 | 0.6 | 0.4 |  |  |  |  |
| Body mass | 1 | 0.9 | 1 | 1 |  |  |  |  |
| SVL | 1 | 0.9 | 0.8 | 0.6 |  |  |  |  |
| Tibiofibula length | 0.9 | 0.9 | 0.6 | 0.4 |  |  |  |  |

Given the limited inferential resolution of the exact tests with three sites per condition, these values were interpreted together with the observed distributions rather than using a conventional significance threshold. Overall, the classic functional indices showed substantial overlap and no consistent separation between contamination conditions. The correlation between functional richness and taxonomic richness was weak (p = 0.35). FRic residuals, which represent relative functional richness after controlling for the effect of species richness, included both positive and negative values in both groups (exact permutation p = 0.5; Table 2), with no consistent directional difference between conditions. Analysis of trait distribution moments likewise revealed substantial variation among sites and broad overlap between contaminated and reference streams for CWM, CWsd, skewness, and kurtosis across the evaluated traits (Table 2). Taking together, these patterns provide no clear evidence of consistent directional filtering or functional homogenization associated with contamination status.

Analysis of trait distribution moments revealed variation among sites, yet no significant differences were detected between contaminated and reference sites for CWM, CWsd, skewness, or kurtosis in any of the evaluated traits (Table 2). Taken together, these results indicate that neither the overall functional space nor the internal distribution of traits within communities showed consistent signals of directional filtering or functional homogenization associated with oil contamination.

Projection of communities onto the multivariate functional space revealed partial overlap between contaminated and reference streams, with no clear separation of centroids along the first two principal axes (Fig. 4). Both global functional dispersion and centroid distance to the overall functional-space center varied among individual sites (Table 3), while the exact permutation p-values for comparisons between conditions were 0.3 for global dispersion and 0.2 for distance to the global centroid. Consistent with the substantial overlap between stream categories, leave-one-out sensitivity analyses indicated that the qualitative pattern was not attributable to any single stream.

**Fig. 4.**
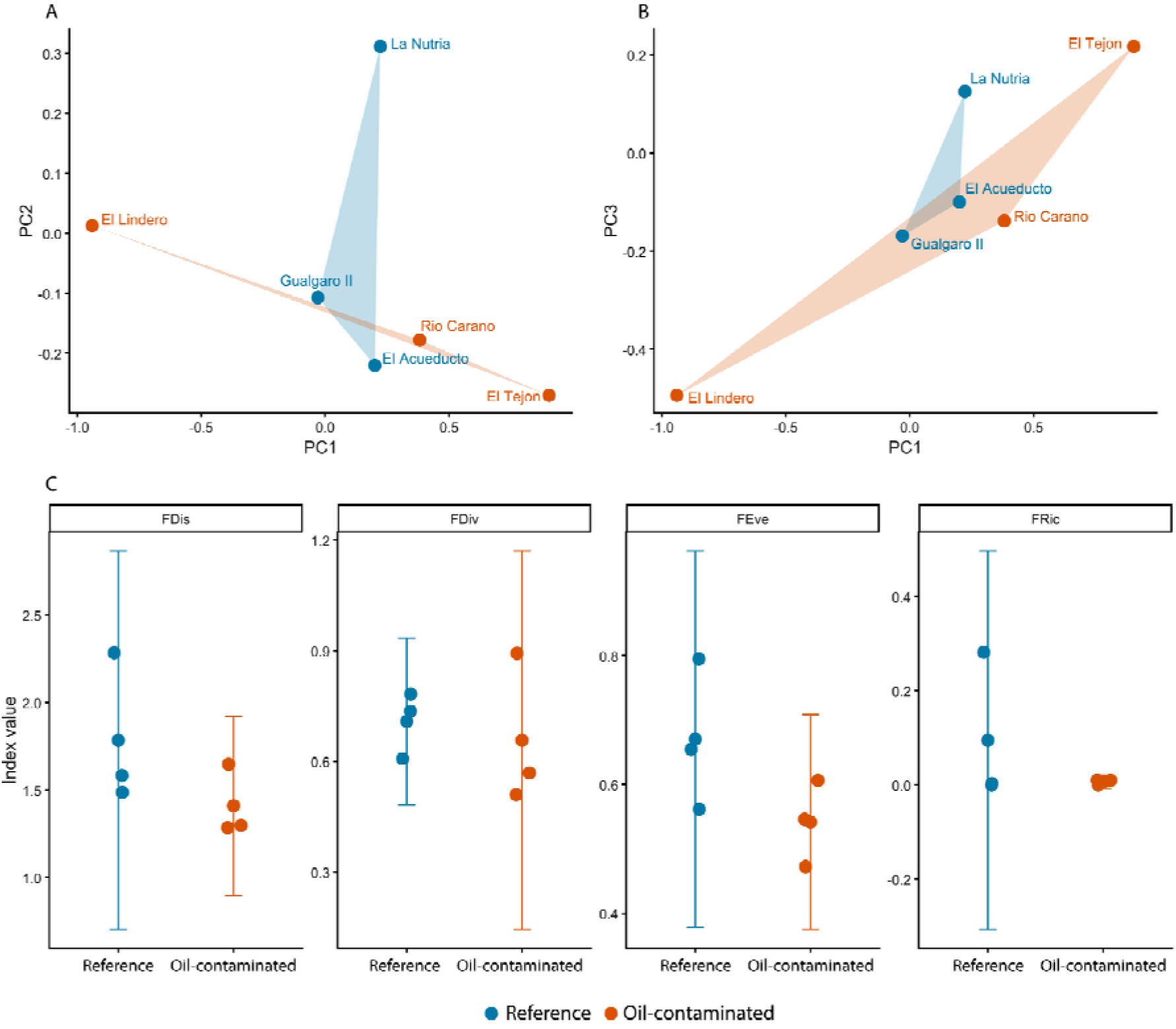
Functional diversity showed broad overlap between oil-contaminated and reference streams, with substantial site-level heterogeneity but no consistent evidence of functional simplification associated with chronic oil contamination. (a–b) Multivariate functional space of anuran assemblages based on principal component axes derived from continuous traits. Points represent stream centroids in trait space and shaded polygons indicate the convex hull occupied by each stream category (reference in blue; oil-contaminated in orange). Assemblages from both stream types showed partial overlap, and differences were driven primarily by variation among individual streams rather than by contamination category. (c) Stream-level values of four global functional diversity indices: functional dispersion (FDis), functional divergence (FDiv), functional evenness (FEve), and functional richness (FRic). Points show individual streams and vertical bars indicate group means ± SD. Functional metrics varied among streams and showed substantial overlap between oil-contaminated and reference streams; exact permutation p-values for each index are reported in Table 2. Together, these analyses indicate that chronic oil contamination was not associated with a clear contraction or reorganization of functional trait space in these frog assemblages

**Table 3.** Multivariate functional structure varied among individual streams but showed no consistent separation between oil-contaminated and reference streams. Stream-level estimates of multivariate functional structure for anuran assemblages sampled in six streams in Barbacoas, Nariño, Colombia. For each stream, we report global functional dispersion (Disp_global), calculated as the abundance-weighted mean distance of species to the community centroid in multivariate trait space, and the distance between each stream centroid and the global centroid (Dist_centroid_global), which reflects the degree of functional displacement relative to the overall regional assemblage. Both metrics varied substantially among individual streams, with broad overlapping between contamination categories

| **Site** | **Site state** | **Disp_global** | **Dist_centroid_global** |
|---|---|---|---|
| Río Caraño | oil-contaminated | 1.3 | 0.381 |
| El Acueducto | reference | 1.58 | 0.261 |
| El Lindero | oil-contaminated | 1.65 | 1.06 |
| El Tejón | oil-contaminated | 1.28 | 0.96 |
| Gualgaro II | reference | 1.49 | 0.147 |
| La Nutria | reference | 2.28 | 0.503 |
| Average categorical functional composition by group |  |  |  |
| Site state | Trait | Cat | Average proportion |
| Reference | Activity | Nocturnal | 1.0 |
|  | Behavior | Perched | 1.0 |
|  | Microhabitat | Arboreal | 0.293 |
|  | Microhabitat | Terrestrial | 0.233 |
|  | Foot webbing | Absent | 0.655 |
| Oil-contaminated | Activity | Nocturnal | 1.0 |
|  | Behavior | Perched | 1.0 |
|  | Microhabitat | Arboreal | 0.35 |
|  | Microhabitat | Terrestrial | 0.3 |

Trait distributions and categorical ecological strategies were broadly similar between oil-contaminated and reference streams (Fig. 5). For continuous traits, including perch height and body mass, there were no consistent shifts in central tendency or trait dispersion between oil-contaminated and reference streams, with variation occurring mainly among individual sites (Fig. 5A; Table 2). Similarly, the abundance-weighted representation of the focal categorical traits showed substantial overlap between stream categories. Microhabitat use, substrate type, and foot webbing varied among individual streams, but none showed a consistent directional separation between reference and oil-contaminated streams (Fig. 5B). Activity period and behavioral state were highly uniform across the dataset, with observations dominated by nocturnal and perched individuals; because these patterns were strongly conditioned by the nocturnal survey protocol, they were treated descriptively and were not interpreted as evidence of functional similarity between stream categories. Exact permutation tests for the focal categorical traits yielded no strong separation between conditions, and leave-one-out sensitivity analyses indicated that the overall qualitative pattern did not depend strongly on any single stream. Collectively, these results provide no clear evidence of consistent functional restructuring associated with contamination status.

**Fig. 5.**
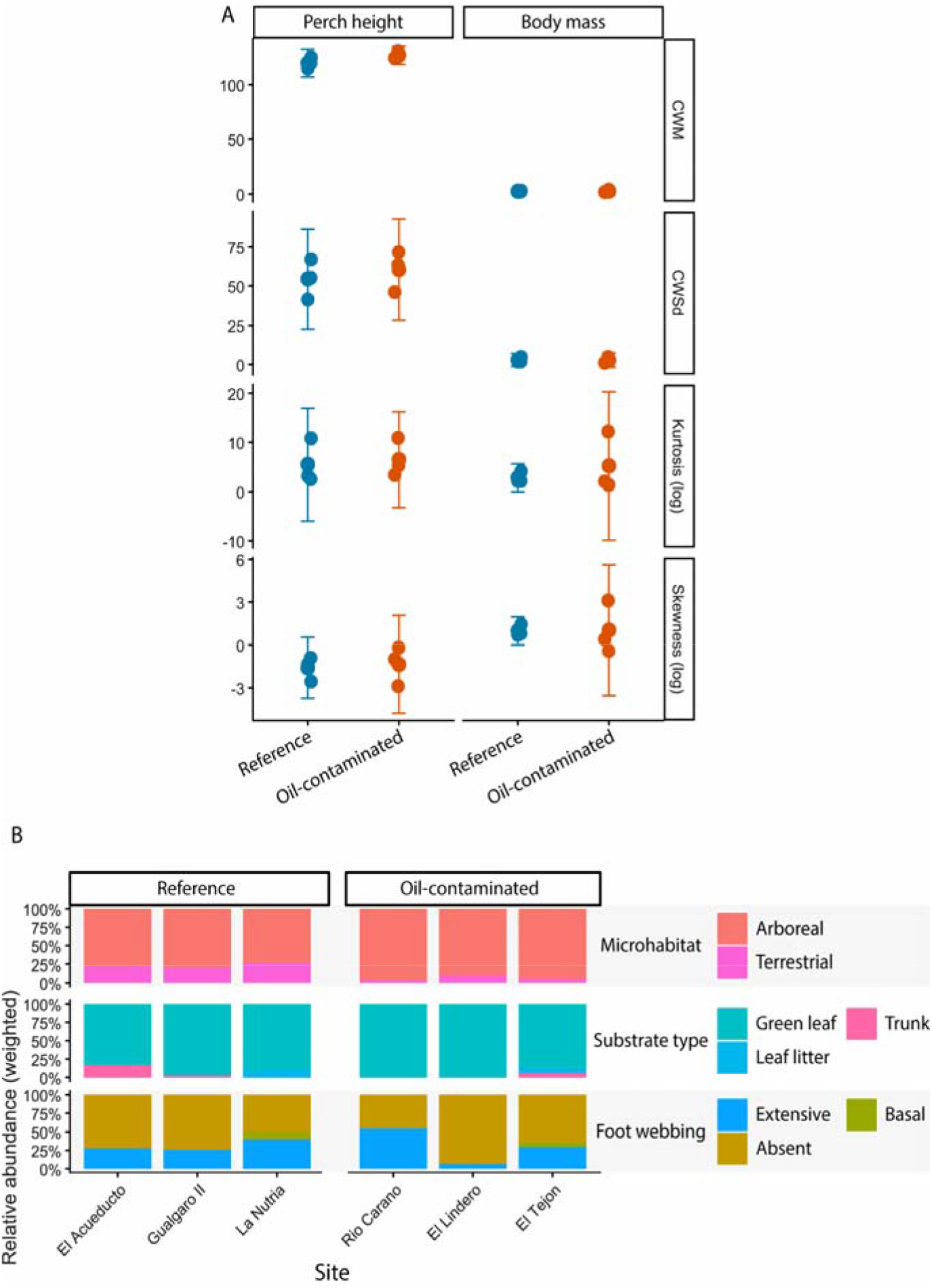
Trait distributions and categorical ecological composition showed substantial overlap between oil-contaminated and reference streams, with variation occurring mainly among individual sites. (a) Distributional moments for representative continuous traits (perch height and body mass), including community-weighted means (CWM), weighted standard deviations (CWsd), kurtosis, and skewness for each stream category. Points represent individual streams and vertical bars indicate group means ± SD. These examples illustrate the broader pattern observed across all measured continuous traits: substantial among-stream variation and broad overlap between stream categories (Table 2). (b) Relative abundance-weighted representation of the focal categorical functional traits across streams: microhabitat use, substrate type, and foot webbing development. These traits varied among individual streams but showed no consistent directional separation between reference and oil-contaminated streams. Activity period and behavioral state are not shown because their observed distributions were strongly constrained by the nocturnal survey protocol and were retained only as descriptive variables. Together, these patterns provide no clear evidence of consistent functional reorganization associated with contamination status

## Discussion

Our results reveal a notable discrepancy between the taxonomic and functional responses of amphibian communities in the Pacific piedmont of Nariño to chronic oil contamination. At the taxonomic level, reference streams collectively contained more species and individuals tended to show higher coverage-standardized diversity than oil-contaminated streams, although substantial variation among individual streams and broad overlap in confidence intervals indicate that this taxonomic pattern was not uniform across sites. Yet this taxonomic signal did not translate into a clear functional restructuring. Across all functional approaches examined, including classical diversity indices, trait distribution moments, multivariate functional space structure, and categorical functional composition, no consistent differences were detected between oil-contaminated and reference streams. Taken together, these results indicate a tendency toward lower taxonomic diversity in streams classified as oil-contaminated, without a corresponding detectable functional simplification at the spatial and temporal scales evaluated here.

This apparent functional stability despite taxonomic change is consistent with expectations for tropical communities characterized by high functional redundancy, where the loss or replacement of individual species can alter composition without producing proportional changes in aggregate functional structure (McGill et al., 2006; Flynn et al., 2009; Mori et al., 2013). This pattern deviates from the classical expectation that chronic pollution acts as an environmental filter that contracts functional space or favors trait convergence (Dolédec & Statzner, 2010; Spasojevic & Suding, 2012; Mouillot et al., 2013). Instead, our results suggest that these communities retain functionally similar species combinations across conditions, maintaining comparable functional metrics despite the overall tendency toward lower species richness and abundance at oil-contaminated sites. Two non-exclusive mechanisms could explain this resilience. First, the exceptional species richness of the Chocó Biogeographic Region increases the probability that multiple species share similar functional roles, providing a form of ecological insurance against disturbance-driven species loss (Naeem, 1998; Yachi & Loreau, 1999), a pattern documented in other species-rich amphibian communities in tropical streams (Strauß et al., 2010). Second, the spatially heterogeneous nature of oil contamination, where spill intensity and persistence vary among sites and microhabitats, may allow the persistence of refugia that sustain functionally diverse assemblages at the local scale (Heard et al., 2015).

The lack of a consistent functional separation between stream categories was evident across all four complementary approaches. Classic diversity indices (FRic, FEve, FDiv, and FDis) showed substantial overlap between oil-contaminated and reference streams, and functional space breadth was not consistently lower in streams classified as oil-contaminated.. Although the reference stream La Nutria occupied a broader and more dispersed functional space than other sites, these differences were not consistent across reference sites as a group. A partial decoupling between taxonomic and functional diversity also reported in other chronically disturbed systems (Mirabel et al., 2020; Ceron et al., 2023), consistent with evidence that in highly diverse systems species loss does not necessarily translate into an immediate loss of ecological function when multiple species occupy similar positions in trait space (Rosenfeld, 2002).

Univariate trait moments told the same story. According to environmental filtering theory, a strong filter should shift mean trait values or reduce trait variability within communities (Shipley, 2009; Spasojevic & Suding, 2012), yet none of the moments examined (CWM, CWsd, skewness, kurtosis) differed between stream types, providing no evidence of a clear directional filtering pattern associated with contamination status in the morphological and ecological traits measured here. Effects may instead act on dimensions not captured by this study, such as physiological tolerance, resistance to chemical stress, reproductive success, or early larval survival, traits known to be highly sensitive to pollutants in amphibians and other ectothermic vertebrates (Rohr & Palme, 2012; Slaby et al., 2019; Martin et al., 2024). At the multivariate level, functional centroids of contaminated and reference streams overlapped broadly. Neither dispersion nor centroid distance from the global center differed significantly, indicating that contamination has not consistently displaced functional composition. This contrasts with systems where marked centroid shifts have been interpreted as functional reorganization (Laliberté & Legendre, 2010; Mouillot et al., 2013), and consistent instead with the use of less-contaminated microhabitats or the dominance of tolerant generalist species (Heard et al., 2015).

Categorical composition provided a similarly cautious picture for traits that could be interpreted independently of survey timing. Microhabitat use, substrate type, and foot webbing showed substantial overlap between stream categories, with variation occurring primarily among individual streams rather than as a consistent directional shift between reference and oil-contaminated sites. In contrast, activity period and behavioral state were not interpreted as independent evidence of functional similarity because all surveys were conducted at night and most detected individuals were perched. Their near-uniform distributions therefore largely reflect the sampling protocol rather than unrestricted community-level variation in these ecological dimensions. This limitation is particularly relevant because diurnal species known from the area, such as *Epipedobates boulengeri* and *E. narinensis*, were not recorded during nocturnal surveys, whereas *Oophaga sylvatica* was detected only sporadically and at a single site. Consequently, our conclusions regarding categorical functional composition are based primarily on microhabitat use, substrate type, and foot webbing rather than on activity period or behavioral state.

Although we did not detect a clear functional restructuring, the observed taxonomic patterns carry important ecological and conservation implications. The lower species richness and abundance recorded at oil-contaminated sites suggest that oil contamination may be associated with reduced components of local biodiversity, even if these losses are not yet reflected in aggregated functional metrics. This is particularly concerning in a region such as the Chocó Biogeographic Region, where exceptional regional diversity is exposed to multiple concurrent anthropogenic pressures, including deforestation, illegal mining, wildlife trafficking, aerial fumigation with herbicides as part of illicit crop eradication programs, and climate change (Portocarrero-Aya et al., 2014; Flórez-Paí, 2016; Negret et al., 2021), compounded by chronic oil contamination associated with the Trans-Andean Pipeline and illegal crude oil extraction and processing (Garcés-Ordóñez & Espinosa, 2019; Ríos-Mármol et al., 2023). In this context, even partial taxonomic losses may have substantial ecological consequences over the medium to long term, particularly if they disproportionately affect rare, endemic, or species with limited dispersal ability or narrow habitat requirements.

The use of multiple complementary functional dimensions, integrating classic indices, trait distribution moments, multivariate functional space, and categorical composition, reduces dependence on any single metric and strengthens the consistency of our conclusions. Similarly, the conservative analytical framework based on exact permutation tests and leave-one-out sensitivity analyses minimizes the risk of detecting spurious patterns given the limited number of sampling sites, a constraint inherent to field research in conflict-affected biodiversity hotspots, where security conditions impose additional restrictions on sampling effort, site access, and working hours that are rarely acknowledged in the literature but represent a persistent challenge for biodiversity science in regions such as the Colombian Pacific (Hanson et al., 2009; Hanson, 2018).

Several limitations should be considered when interpreting these results. The small number of sites inherently limits statistical power to detect subtle differences between conditions. Furthermore, we were unable to reconstruct the temporal history of each oil spill or the actual hydrocarbon concentration or duration of exposure at each site. Such heterogeneity may obscure contrasting biological responses within the oil-contaminated stream category. Similarly, the functional traits analyzed primarily capture general morphological and ecological dimensions and may not reflect sublethal physiological responses to oil-contamination. Finally, although sampling spanned different seasons, the study did not include temporal monitoring of community degradation or post-disturbance recovery trajectories.

In summary, amphibian communities in oil-contaminated streams of the Pacific piedmont of Nariño showed taxonomic patterns that generally tended toward lower richness, abundance, and diversity than those in reference streams, although these differences varied among individual sites and were not accompanied by a clear functional simplification. This decoupling underscores the importance of evaluating taxonomic and functional diversity independently, rather than assuming that both respond in parallel to anthropogenic disturbance. Future studies should integrate long-term monitoring, more precise characterization of hydrocarbon exposure intensity and duration, physiological and demographic traits, and other taxonomic groups occupying aquatic and riparian habitats, to assess whether the functional stability documented here reflects genuine ecological resilience or an early stage of a longer trajectory of functional degradation.

## Acknowledgments

We thank the individuals and institutions that made this study possible. We extend special thanks to the indigenous communities of the Awá Gran Sábalo Indigenous Reserve and the +nkal Awá La Nutria Piman, Bangsias Bird Lodge, and Río Ñambí nature reserves. We are also grateful to Miguel García, *Estiven Pérez (in memoriam)*, Silvana Pahi, Alvaro Pai, Alberto García, Eulicer Pascal, Carlos Manuel Nastacuaz, Jefferson David Guanga, José López, Diego Ruíz, and Rosita Paí for their invaluable support during fieldwork. We thank Dr. Andrew Crawford for his support and assistance in facilitating the required research permits. Finally, we thank Melissa Achicanoy, Yasmin Betancourt, Fernando Santander, and Brandon Buitrago for their expertise on the socio-environmental dimensions of crude oil spills and their assistance with field data collection.

## Statements and Declarations

### Author Contributions

**MBC:** Conceptualization, Methodology, Investigation, Data curation, Writing – original draft, Writing – review & editing, Project administration. **MPT:** Conceptualization, Methodology, Investigation, Formal analysis, Writing – original draft, Writing – review & editing. **JRO:** Formal analysis, Writing – original draft, Writing – review & editing. **CFP:** Resources, Project administration. **MVP:** Investigation. **LO:** Funding acquisition, Writing – review & editing. All authors reviewed and approved the final version of the manuscript.

### Data Availability

The datasets generated during and/or analysed during the current study are available in the Zenodo repository (https://doi.org/10.5281/zenodo.23399649).

